# Assessing *Myf5* and *Lbx1* contribution to carapace development by reproducing their turtle-specific signatures in mouse embryos

**DOI:** 10.1101/2022.01.18.476730

**Authors:** Triin Tekko, Ana Nóvoa, Moisés Mallo

## Abstract

**Background:** The turtle carapace is an evolutionary novelty resulting from changes in the processes that build ribs and their associated muscles in most tetrapod species. Turtle embryos have several unique features that might play a role in this process, including the carapacial ridge, a *Myf5* gene with shorter coding region that generates an alternative splice variant lacking exon 2, and unusual expression patterns of *Lbx1* and *HGF*.

**Results:** We generated *Myf5* alleles reproducing the *Myf5* turtle expression features. At mid gestation, mouse embryos expressing *Myf5* lacking exon 2 reproduced some early properties of turtle somites, but still developed into viable and fertile mice. Extending *Lbx1* expression into the hypaxial dermomyotomal lip of trunk somites to mimic the turtle *Lbx1* expression pattern, produced fusions in the distal part of the ribs.

**Conclusions:** Turtle-like *Myf5* activity might generate a plastic state in developing trunk somites under which they can either enter carapace morphogenetic routes, possibly triggered by signals from the carapacial ridge, or still engage in the development of a standard tetrapod ribcage in the absence of those signals. In addition, trunk *Lbx1* expression might play a later role in the formation of the lateral border of the carapace.

## INTRODUCTION

The turtle carapace represents one of the most drastic deviations from the standard tetrapod body plan. Formation of the carapace involves a combination of various processes that change the growth and differentiation properties of the rib-forming area of the skeleton^1,2^. Ribs develop from the sclerotome of trunk somites resulting from inductive interactions from the adjacent myotome ^3–10^. In most tetrapod species these somites extend ventro-distally along the body wall to surround the coelom, eventually generating the rib cage and its associated muscles. In the turtle embryo, however, trunk somites undergo what has been called “axial arrest” ^2^. This entails blocking somite ventro-distal growth around the stage when they enter abaxial development ^11^. As a result, the ventral lips of the somites fail to extend further distally and grow laterally towards the carapacial ridge (CR), a longitudinal crest along the lateral aspect of the body wall ^1,12–14^. It is thought that the CR then regulates different events during differentiation of the distal end of these somites, including their growth and differentiation along the anterior-posterior axis, eventually covering dorsally the developing limb buds ^2^. In addition to the distinct growth characteristics of the distal somitic borders, their myotome also undergoes an alternative differentiation route, producing a fibrous mass between the rib anlages instead of the muscles normally produced in other tetrapods ^1,15^.

The molecular bases for the divergent somite development leading to carapace formation are far from being clarified. A series of molecular and expression studies have identified several factors that could play relevant roles in this process. It has been shown that in turtles the ventro-lateral (hypaxial) lip of the dermomyotome undergoes premature de-epithelization, which could contribute to “axial arrest” and to the thinning of the myotome observed in turtle embryos ^14^. The abnormal behavior of the hypaxial dermomyotomal lip of trunk somites has been linked to the unusual expression patterns of *Lbx1* and *HGF*, two factors normally involved in the formation of limb muscle precursors from limb somites ^16–19^. Normally in tetrapod embryos, *Lbx1* expression is restricted to the hypaxial dermomyotomal lip at cervical and limb regions ^16^ and HGF is expressed in the developing limb buds ^17,18^. In turtle embryos, their expression is extended to the interlimb area, where *Lbx1* is activated in the hypaxial dermomyotomal lip, while *HGF* shows a dynamic pattern involving the sclerotome and the lateral embryonic body wall ^14^. Given the role that these two factors play in the behavior of cells in the hypaxial dermomyotomal lip, it has been suggested that they might also take part in the accelerated differentiation of the hypaxial dermomyotomal lip of the trunk somites in turtles ^14^. Interestingly, blocking HGF activity using a molecular inhibitor resulted in the loss of the CR, suggesting that this growth factor plays a role in the induction of this structure ^14^.

Distinct turtle-specific sequence and expression characteristics of the *Myf5* gene could also play a role in the divergent differentiation routes of their somites ^20^. It has been described that the Myf5 proteins of turtles contain a four amino acid deletion in the activating domain of the proteins, corresponding to the 5’ end of exon 3 ^20^. In addition, molecular analyses in *Pelodiscus sinensis* embryos revealed the presence of an alternatively spliced transcript that skips exon 2, leading to a frame shift in the open reading frame that removes part of the activating domain ^20^. Such alternative transcript has never been described in any other tetrapod. Given the instrumental role of Myf5 in the myotomal-sclerotomal interactions leading to rib formation ^4,6,8,10,21^, it has been suggested that this gene could have played an important role in the emergence of the carapace ^14^. Whether this is indeed the case remains to be determined.

In this work we assessed the possible role of the two turtle-specific features of the *Myf5* gene on the development of the carapace using genetic approaches in mice. We first generated three mouse strains, each containing a *Myf5* allele with turtle-specific characteristics: an allele containing the 4 amino acid deletion, an allele lacking exon 2, thus forcing the production of transcripts mimicking the turtle splice variant, and a third allele combining the features of the other two alleles. We found that alleles lacking exon 2 influence the differentiation of the dermomyotome and affect gene expression in both the myotome and dermomyotome. Interestingly, despite these molecular findings, mice carrying either of these alleles in homozygosity were viable and fertile, showing fully penetrant, but rather mild skeletal phenotypes involving the first two ribs. We also assessed the functional relevance of *Lbx1* expression in the hypaxial dermomyotomal lip of trunk somites by generating transgenic embryos reproducing this turtle-specific feature. These transgenic mice exhibited distal rib fusions, suggesting a possible role for trunk *Lbx1* expression in the formation of the lateral border of the carapace.

## RESULTS

To evaluate the influence of *Myf5* on the generation of the turtle body plan we generated mouse models containing a *Myf5* locus resembling the characteristics of the turtle gene. The *Myf5* exon 3 of turtles lacks 12 nucleotides from the 5’ end of exon 3 when compared to other tetrapods ^20^, resulting in a four amino acid deletion within the protein’s activation domain ^20,22^. Comparison of the intron 2-exon 3 boundary between mouse and turtle (*P. sinensis*) shows that the turtle gene lost the splice acceptor normally used in the mouse and reconverted a downstream AG into the exon3 splice acceptor (Fig. 1). To generate a mouse *Myf5* allele reproducing this feature of the *Myf5* turtle gene (*Myf5^Tr-ex3^*) we removed the first 12 nucleotides of the wild type exon3, keeping the structure of the acceptor site normally used in mice. Mice homozygous for this allele were viable and fertile, showing no apparent phenotypic alteration. *Myf5* expression from this allele resembled that of the normal mouse *Myf5* gene according to *in situ* hybridization staining (Fig. 1). Skeletal analyses of E18.5 *Myf5^Tr-ex3/Tr-ex3^* fetuses also failed to show any effect on rib development (n=8) (Fig. 2).

**Figure 1.**
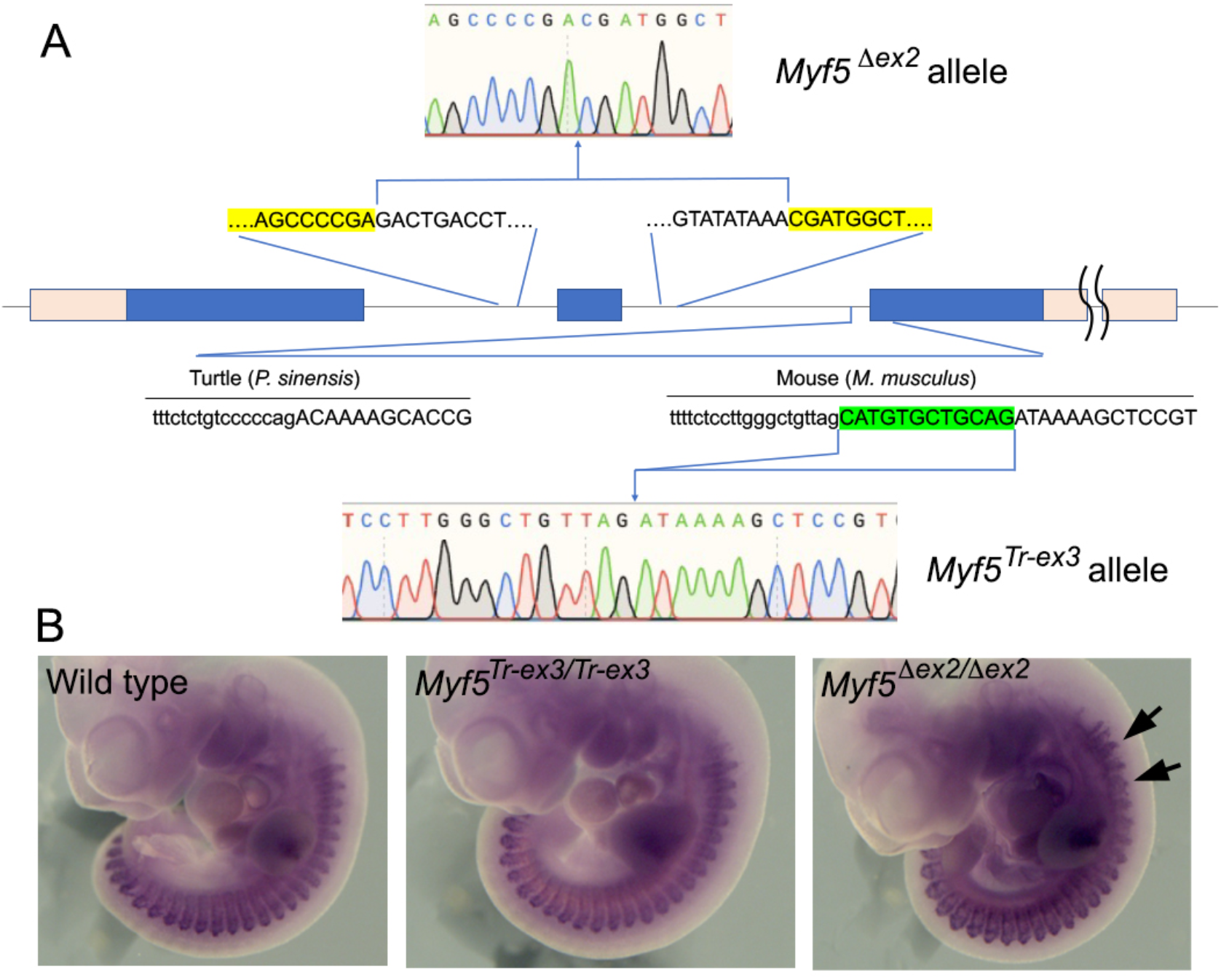
Generation of the mutant *Myf5* alleles. A. Schematic representation of the *Myf5* gene. The blue boxes represent the coding region encoded by three exons; the untranslated regions of the gene are represented as pink boxes and the introns as lines between the exons. The position within the gene and the sequences of the *Myf5^Δex2^* and *Myf5^Tr-ex3^* alleles are shown. Highlighted in yellow are the borders of the sequence deleted in the *Myf5^Δex2^* allele and in green the 12 nucleotide region deleted in *Myf5^Tr-ex3^*. Also shown is the intron-exon boundary of the turtle *P. sinensis Myf5* gene. B. Whole mount *in situ* hybridization for *Myf5* in E10.5 wild type and homozygous embryos for the *Myf5^Tr-ex3^* and *Myf5^Δex2^* alleles. Arrows indicate some altered morphology of the *Myf5* signal in rostral somites.

**Figure 2.**
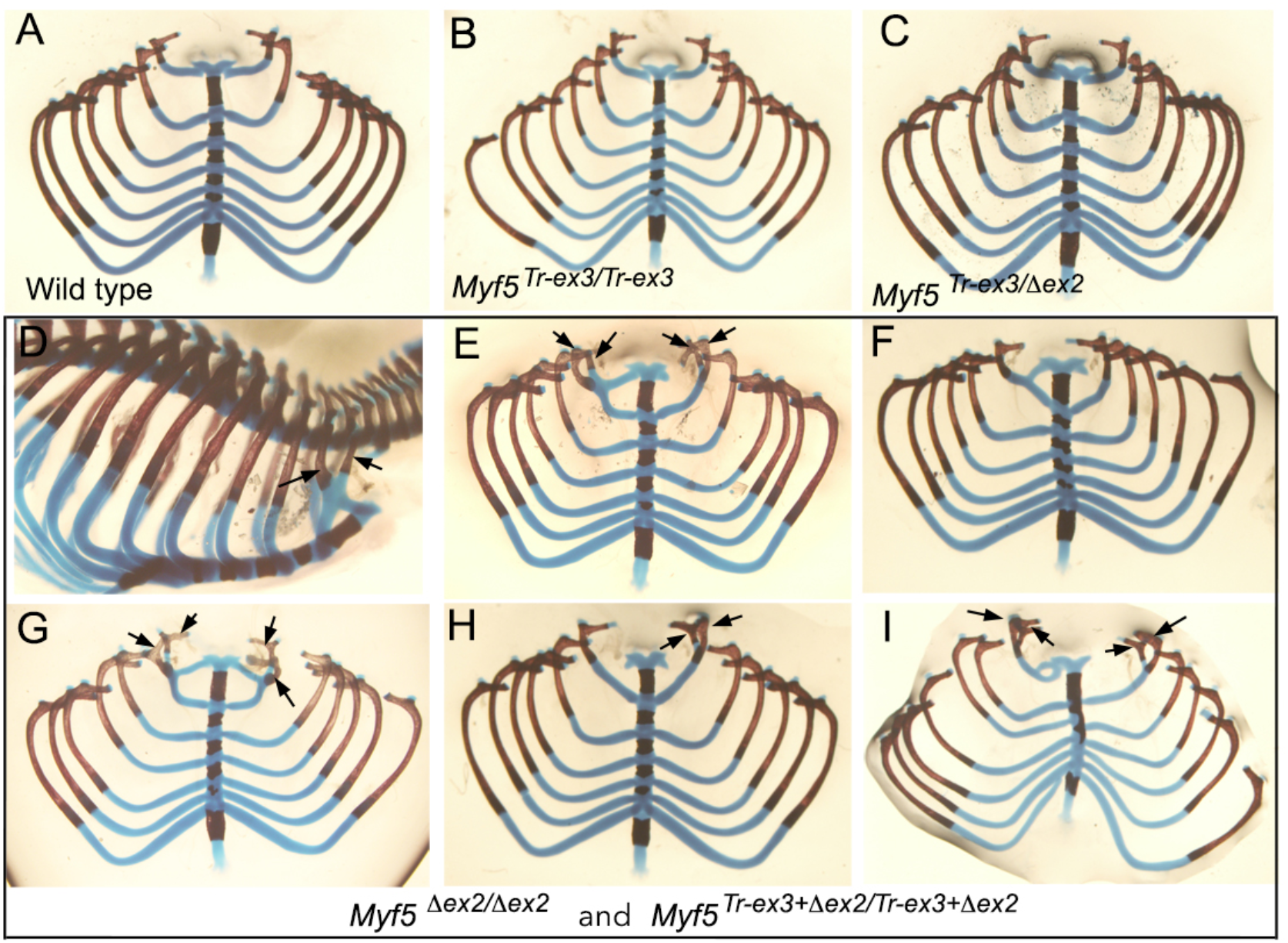
Dissected rib cages of E18.5 fetuses with different combinations of *Myf5* alleles. A. Wild type control. B. Homozygous mutant for the *Myf5^Tr-ex3^* allele. C. Trans-heterozygote mutant for the *Myf5^Tr-ex3^* and *Myf5^Δex2^* alleles. D-I. Examples of the phenotypes observed in homozygous mutants for both *Myf5^Δex2^* and *Myf5^Tr-ex3+Δex2^*. The pattern shown in I was obtained only once in a *Myf5^Δex2^* homozygous mutant. The arrows indicate the proximal part of ribs 1 and 2 when fused ventrally, best illustrated in the specimen shown in D before dissection.

To mimic the second turtle-specific characteristic of *Myf5* expression, namely the production of an alternatively spliced transcript lacking exon 2, we generated a second *Myf5* allele lacking this exon (*Myf5^Δex2^*). This allele also produced transcripts with expression patterns resembling those observed for *Myf5* in wild type embryos, although it is possible that expression in the hypaxial region of the myotome was slightly stronger than in control embryos (Fig. 1). In addition, some *Myf5^Δex2/ Δex2^* embryos had subtle alterations in the shape of the of the *Myf5* signal in the most rostral somites. *Myf5^Δex2^* homozygous mice were also viable and fertile. However, skeletal analyses of E18.5 *Myf5^Δex2/ Δex2^* fetuses revealed morphological alterations affecting the first two ribs. Phenotypic alterations were observed in all (n=15) *Myf5^Δex2/ Δex2^* fetuses analyzed, although their patterns were slightly variable, including various types of fusions between the first and second ribs, both unilaterally or bilaterally, and in some cases also total absence of the first rib (Fig. 2). We could not detect any other alteration in the skeletons of these fetuses. None of these features were observed in *Myf5^Δex2^* heterozygotes (n=11).

To assess possible interactions between both types of turtle *Myf5* transcripts, we generated trans-heterozygous mice for the *Myf5^Tr-ex3^* and *Myf5^Δex2^* alleles. *Myf5^Tr-ex3/ Δex2^* animals were also viable and fertile, showing no detectable abnormal phenotypic trait, including a normal rib cage (n=14) (Fig. 2).

Finally, we generated a third *Myf5* allele containing both absent exon 2 and the shorter exon 3 (*Myf5 ^Δex2/ Tr-ex3^*). This allele would produce transcripts more closely resembling the structure of the alternative spliced transcript of turtles. Homozygous mice for this allele were also viable and fertile and presented fully penetrant skeletal phenotypes (n=17) undistinguishable from those observed in *Myf5 ^Δex2 Δex2^* mutants.

### Molecular characterization of *Myf5* mutant embryos

Given the alterations observed in the first two ribs, we evaluated how the *Myf5* mutation affected different aspects of somite development. We could not identify major changes in the expression patterns of sclerotomal markers in homozygous embryos for either *Myf5 ^Δex2^* or *Myf5 ^Δex2+Tr-ex3^* that could be associated with their rib phenotypes (Fig. 3). However, alterations were readily identified in the myotome and dermomyotome of these mutant embryos. As those alterations were similar in both *Myf5 ^Δex2/ Δex2^* and *Myf5 ^Δex2+Tr-ex3/ Δex2+Tr-ex3^* embryos (most likely because the phenotype depends exclusively on the absence of exon 2), for simplicity, we will refer to them collectively as *Myf5^turtle^* embryos throughout the manuscript.

**Figure 3.**
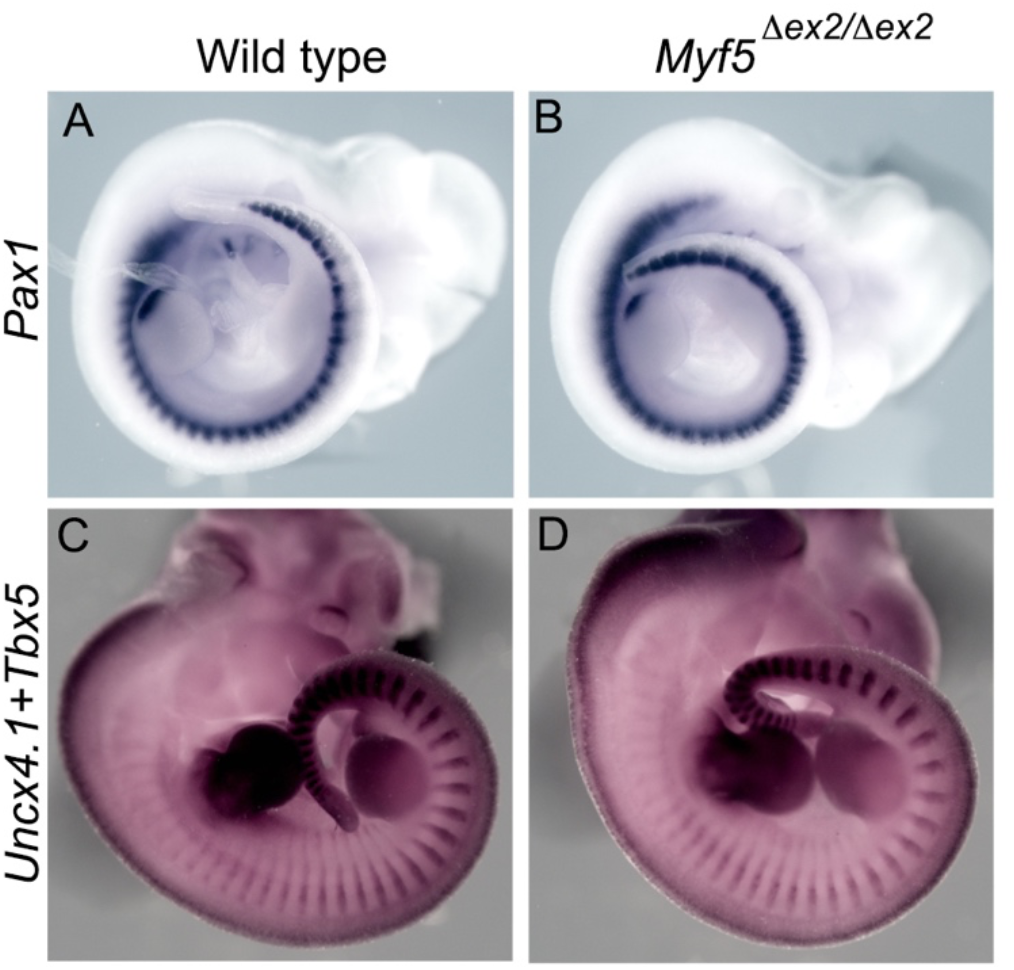
The sclerotome is not affected in homozygous mice for the *Myf5^turtle^* alleles. Whole mount *in situ* hybridization for *Pax1* (A, B) or *Tbx5* and *Uncx4.1* (C, D) in wild type (A, C) and *Myf5^Δex2/ Δex2^* (B, D) embryos at E10.5 (A, B) and E11.5 (C, D).

*Myf6* expression was among the most clearly affected features in *Myf5^turtle^* mutants. In wild type embryos, this gene is activated following both rostral to caudal and epaxial to hypaxial sequences, starting in the epaxial myotome of rostral somites around E9.0 (23 somite pairs) ^23^. In *Myf5^turtle^* embryos, *Myf6* was prematurely activated in the hypaxial border of trunk somites, already clearly visible in embryos with 23 somite pairs, when it was still undetectable in wild type control embryos (Fig. 4). Conversely, at this stage epaxial *Myf6* expression in the more rostral somites was reduced when compared to control embryos. Surprisingly, at E10.5 *Myf6* expression in trunk somites was mostly downregulated in the medial and hypaxial regions of the myotome in *Myf5^turtle^* embryos, a stage when this gene was readily detected throughout the myotome in control embryos (Fig. 4). These observations indicate that the changes introduced in the *Myf5* locus to generate the *Myf5 ^Δex2^* or the *Myf5 ^Δex2/ Tr-ex3^* alleles impacted the regulation of *Myf6* expression, most particularly in the hypaxial myotome. The myotome of *Myf5^turtle^* embryos also showed a significantly reduced *Myogenin* (*Myog*) expression (Fig. 4), affecting both the central region of trunk somites and the central and hypaxial parts of more rostral somites. We could not observe any of these molecular phenotypes in *Myf5^Tr-ex3/Tr-ex3^* or in trans-heterozygotes for the *Myf5 ^Δex2^* and the *Myf5 ^Tr-ex3^* alleles.

**Figure 4.**
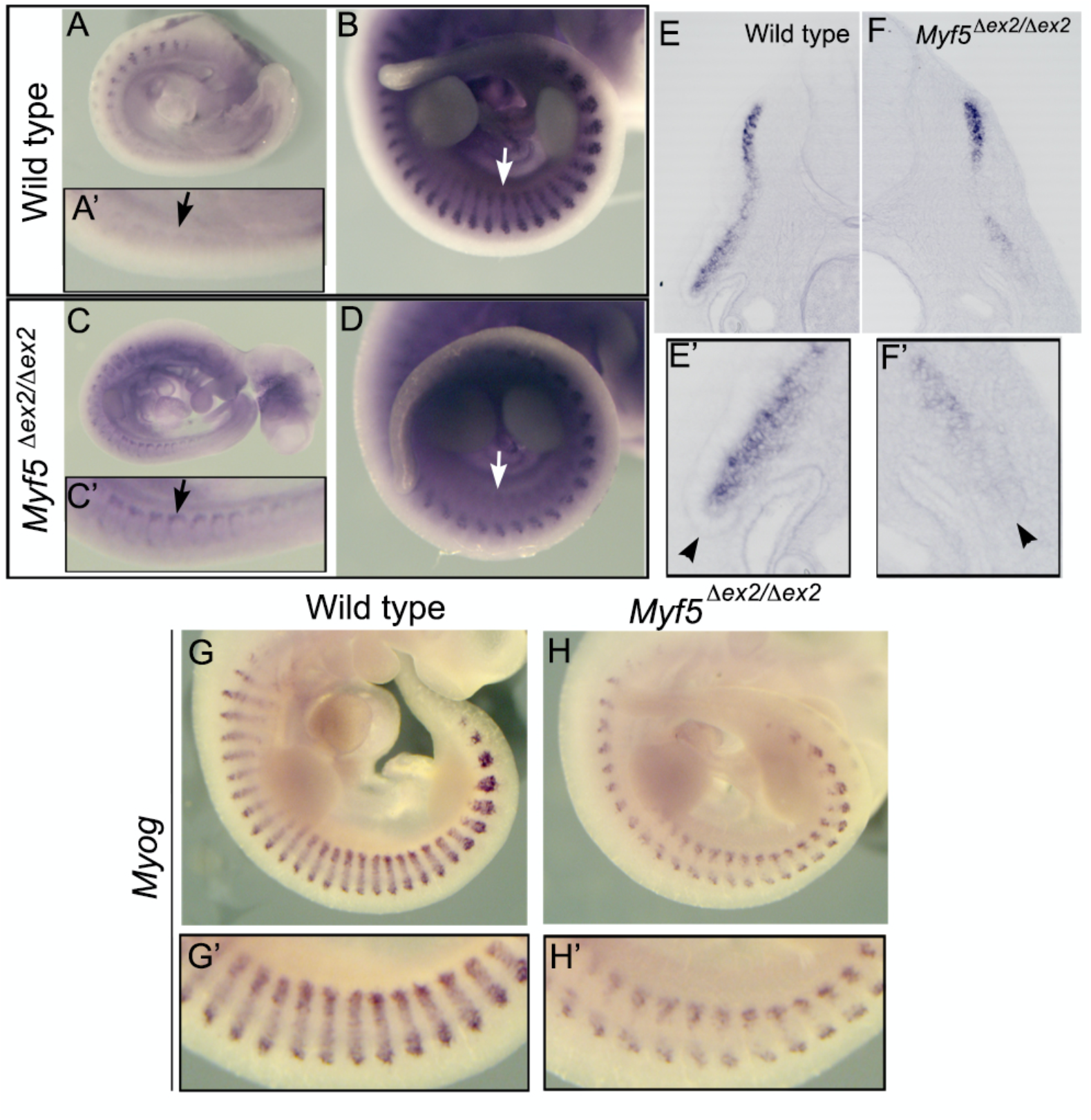
The myotome is affected in homozygous mice for the *Myf5^turtle^* alleles. Whole mount *in situ* hybridization for *Myf6* (A-F’) and *Myog* (G-H’) in wild type (A, A’, B, E, E’, G, G’) and *Myf5^Δex2/ Δex2^* (C, C’, D, F, F’, H, H’) embryos at E9.0 (23 somites) (A, A’, C, C’) and E10.5 (B, D, E-H). Arrows in A’ and C’ indicate premature *Myf6* activation in the hypaxial dermomyotomal lip of *Myf5^Δex2/ Δex2^* embryos. Arrows in B and D indicate the hypaxial myotome of trunk somites. E-F’ show transverse sections through trunk somites of E10.5 embryos. Arrowheads point at the hypaxial dermomyotomal lip.

A close analysis of sections through somites of *Myf6*-stained E10.5 embryos, in addition to the reduction of hypaxial *Myf6* expression, showed an apparent alteration in the morphology of the dermomyotomal hypaxial lip (Fig. 4). We confirmed the abnormal morphology of this region in *Myf5^turtle^* embryos through *Pax3* expression analysis. In wild type embryos, this gene labels the dermomyotome, showing stronger expression in its epithelial hypaxial lip ^24^ (Fig. 5). In *Myf5^turtle^* embryos *Pax3* was also expressed throughout the dermomyotome, but its hypaxial pattern was slightly different to that observed in control embryos (Fig. 5). Sections through this area confirmed the abnormal morphology of the hypaxial dermomyotomal lip in *Myf5^turtle^* embryos, which showed less epithelial morphology than in wild type controls. This pattern somehow resembles the premature de-epithelization of the hypaxial lip of the dermomyotome that has been described as one of the hallmarks of turtle trunk somites ^14^.

**Figure 5.**
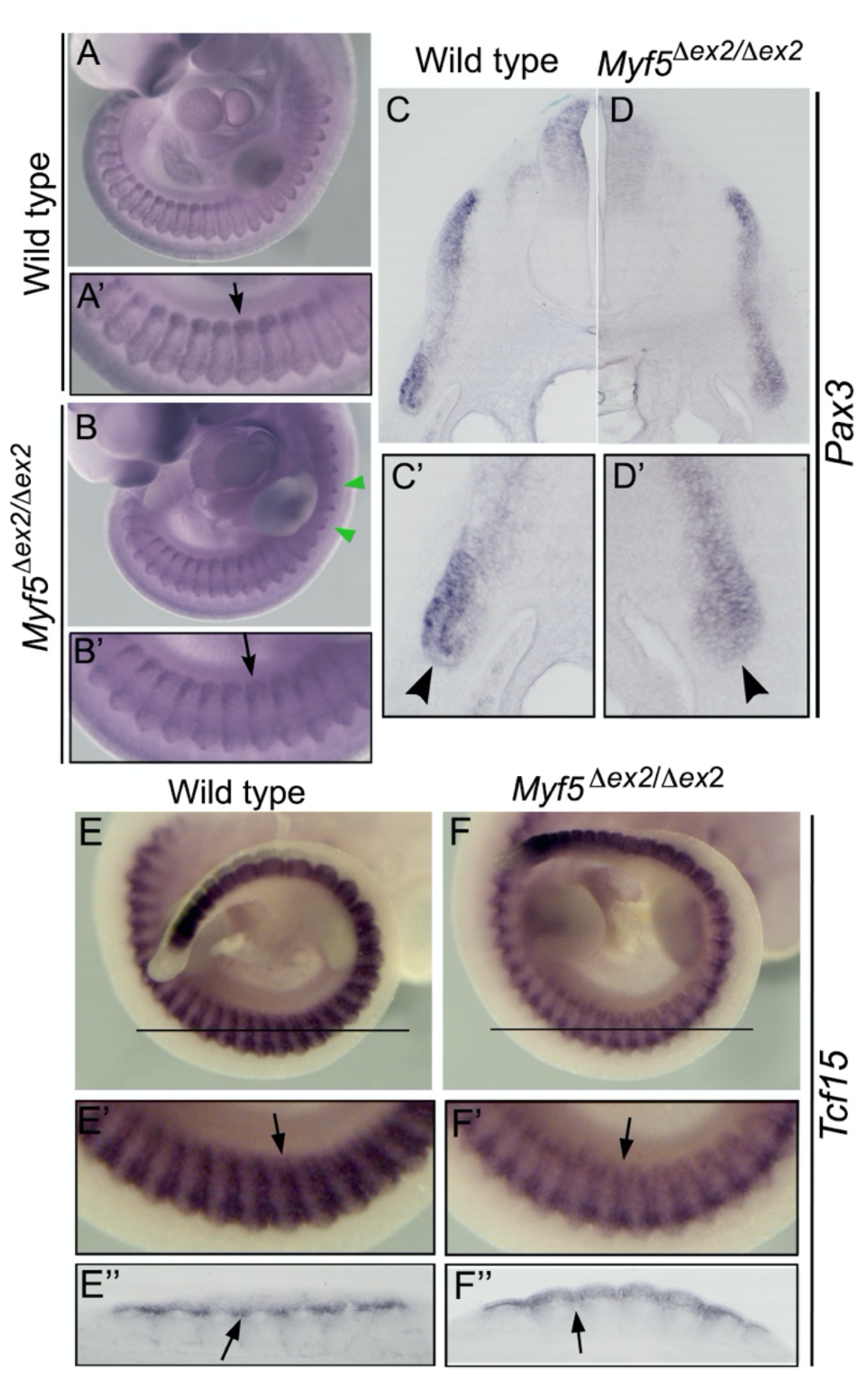
The dermomyotome is affected in *Myf5^turtle^* homozygous embryos. E10.5 wild type (A, A’, C, C’, E, E’, E’’) and *Myf5^Δex2/ Δex2^* (B, B’, D, D’, F, F’, F’’) embryos were stained for *Pax3* (A-D) or *Tcf15* (E, F) by whole mount *in situ* hybridization. Green arrowheads in B indicate rostral somites with morphology change in the mutant embryo; arrows in A’ and B’ indicate the hypaxial dermomyotomal lip of a trunk somite. C-D’ show transverse sections through a trunk somite. The arrowheads in C’ and D’ indicate the different morphologies of the hypaxial dermomyotomal lip in wild type and *Myf5^Δex2/ Δex2^* embryos. E’’ and F’’ show frontal sections through the area indicated in E and F. Arrows in E’, E’’, F’ and F’’ indicate the position of the lateral (rostral and caudal) lips of the dermomyotome.

Alterations in the dermomyotome were not restricted to the hypaxial lip but were also observed in the anterior and posterior lips of the somites as shown by *Paraxis* (*Tcf15*) expression (Fig. 5).

Remarkably, despite the differences observed in the dermomyotome and myotomes of *Myf5^turtle^* embryos, we could not observe any major difference in muscle anatomy in E18.5 fetuses, which also fits with the apparent absence of phenotypes in the adult mutant animals (Fig. 6).

**Figure 6.**
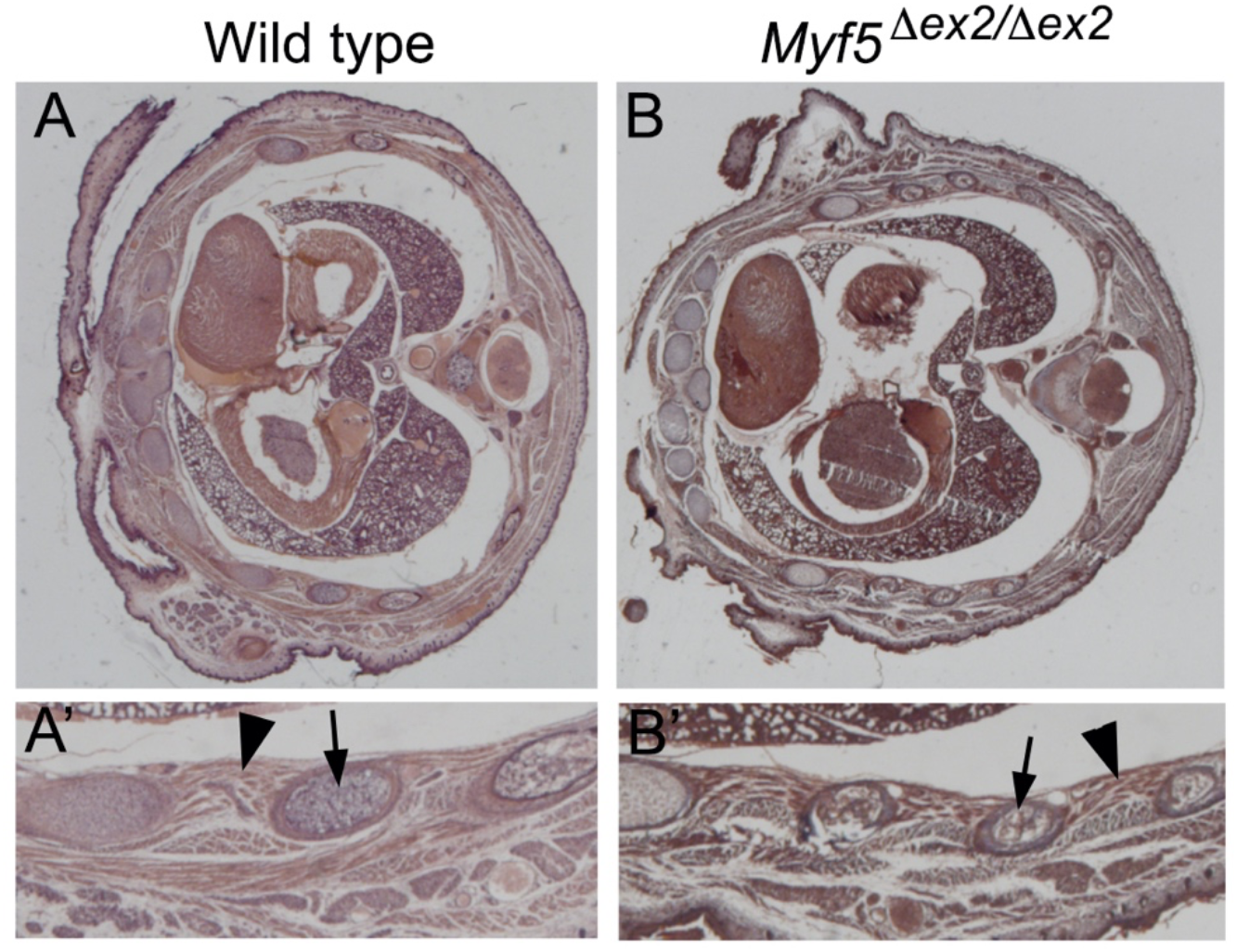
Histological analysis of wild type and *Myf5^turtle^* fetuses. Transverse sections through the thoracic region of E18.5 wild type and *Myf5^Tr-ex3+Δex2/Tr-ex3+Δex2^* fetuses stained with Masson’s trichrome. A’ and B’ show close ups of the ribs (arrows) and their associated muscles (arrowheads).

These results indicate that *Myf5* can indeed introduce some changes in somite development resembling those occurring during the early stages of carapace formation but that such changes are not themselves sufficient to generate any of the features of the mature carapace.

### Activating *Lbx1* in the hypaxial dermomyotomal lip

A distinct characteristic of turtle embryos is the extended somitic *Lbx1* expression into the interlimb area ^14^. *Lbx1* has been shown to stimulate de-epithelization of the dermomyotomal lip to promote migration of the limb muscle progenitors ^16,19^. Because *Myf5^turtle^* embryos also showed loss of epithelial characteristics of the dermomyotomal lip of trunk somites, we tested whether this could be related to upregulated *Lbx1* expression in this embryonic region. We could not detect any significant change in *Lbx1* activation in these mutant embryos (Fig. 7), thus suggesting that *Lbx1* activation in the interlimb area of turtle embryos might not depend on Myf5 activity.

**Figure 7.**
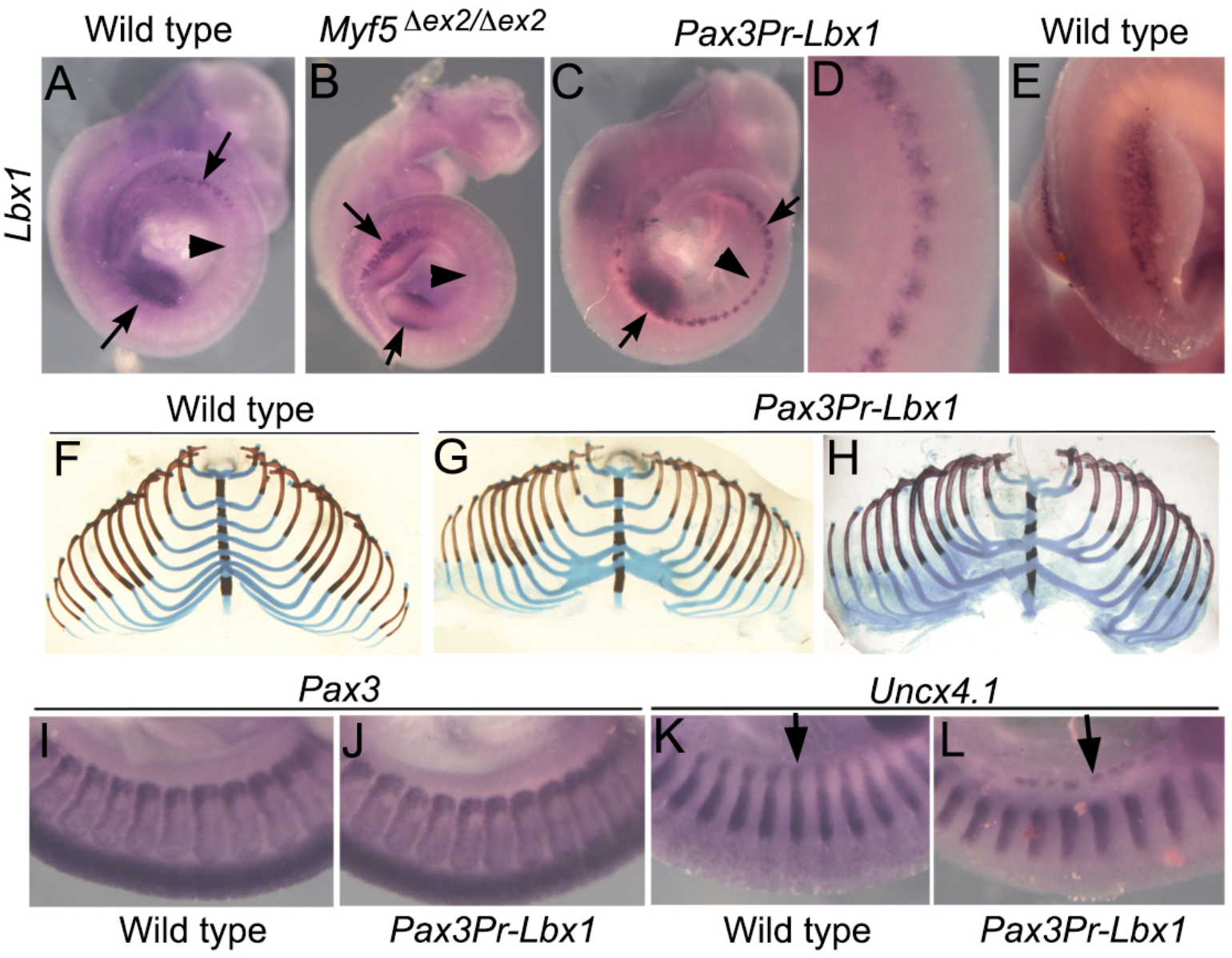
*Lbx1* expression in interlimb somites. A-E. *Lbx1* expression in E10.5 wild type (A, E), *Myf5^Δex2/Δex2^* (B) and *Pax3Pr-Lbx1* transgenic (C, D) embryos. Arrows indicate expression at limb levels and arrowheads indicate the interlimb area. D shows a close up of the interlimb area of the *Pax3Pr-Lbx1* transgenic to illustrate the absence of migratory activity of *Lbx1*-expressing cells. E shows a close up of the hindlimb of a wild type embryo to illustrate the migration of *Lbx1*-positive cells into the limb bud. F-H. Dissected rib cages of E18.5 wild type and *Pax3Pr-Lbx1* transgenic fetuses to illustrate two examples of distal rib fusions in the transgenics. I-L. Close ups of interlimb somites of E10.5 wild type (I, K) and *Pax3Pr-Lbx1* transgenic (J, L) embryos stained for *Pax3* (I, J) or *Uncx4.1* (K, L). The arrows indicate the distal end of the developing somite.

We then tested the possible role of *Lbx1* expression in the interlimb hypaxial dermomyotome in carapace development. For this, we generated transgenic fetuses (*Pax3Pr-Lbx1*) expressing *Lbx1* under the control of the *Pax3* enhancer that activates expression specifically in the hypaxial dermomyotomal lip ^25^ (Fig. 7). From the 13 transgenic fetuses that we collected at E18.5, 8 exhibited various patterns of rib fusions in their distal, cartilaginous region (Fig. 7). These observations were somewhat surprising considering that *Lbx1* has been shown to be involved in limb and diaphragm muscle precursor migration from the hypaxial dermomyotome without obvious effects on skeletal development ^16,19^. We therefore analyzed somite development in *Pax3Pr-Lbx1* embryos at E10.5. Analysis of *Lbx1* expression in these transgenic embryos showed that, contrary to what is observed at limb levels, where *Lbx1*-positive cells migrate into the limb buds, we did not observe any migratory activity in *Lbx1*-positive cells in the interlimb area (Fig. 7). *Pax3* expression in *Pax3Pr-Lbx1* transgenic embryos was also not affected, indicating that at least under these conditions *Lbx1* does not interfere the epithelial characteristics of the hypaxial dermomyotomal lip, and that these cells had not gained migratory properties. *Uncx4.1* expression was, however, affected by *Lbx1* expression in the hypaxial dermomyotomal lip of trunk somites. We observed a disruption of *Uncx4.1* expression in the distal part of the developing somite, where it lost the normal labeling of the posterior compartment, presenting instead a disorganized pattern, detached from the proximal somite region, and reduced in intensity (Fig. 7). We found this pattern in the three transgenic embryos analyzed, indicating that it is a bona fide effect of the ectopic *Lbx1* expression. These observations suggest that *Lbx1* expression in the hypaxial dermomyotomal lip of trunk somites interferes with the normal interactions between the myotome/dermomyotome and the sclerotome involved in rib formation.

## DISCUSSION

In this work we relied on mouse genetics to evaluate the contribution of *Myf5* to carapace formation by generating mice containing alleles resembling turtle *Myf5*-specific characteristics. The *Myf5^turtle^* embryos seemed to reproduce some of the early features observed in the somites of turtle embryos, including the premature de-epithelialization of the hypaxial dermomyotomal lip of trunk somites and deficient myotome development ^14,26^. It is therefore possible that Myf5 activity, and most particularly the activity provided by the splice variant lacking exon 2, might indeed play a role during early steps of carapace development, although our data also show that the activity provided by turtle type Myf5 proteins is clearly insufficient to generate any of the characteristic features of the mature carapace. What role would then *Myf5* play in this process? One possibility is that this gene is required for initial steps of carapace development preparing trunk somites to respond to CR-derived signals that lead to the axial arrest of rib development and their lateral growth that mark the beginning of carapace morphogenesis. In the absence of activities provided by the CR, somites would resume their normal developmental program, eventually rescuing the dermomyotome and myotome alterations to build a standard tetrapod ribcage. Indeed, *Myf5^turtle^* embryos generate muscle masses with apparent normal distribution and function despite the myotome deficiencies observed at mid-gestation. Cell tracing analyses showed that adult muscles derive from a second wave of myotomal cells that replace the primary myotome produced at early developmental stages ^27,28^. It is possible that the normal adult muscles of *Myf5^turtle^* embryos result from the recovery of the deficient myotome early in development by the second myotomal wave. This hypothesis also suggests a possible mechanism for the distinctive differentiation of the turtle myotome: if axial arrest interferes with the activation of the second wave of myotomal cells, the primary myotome might be unable to produce the normal set of rib-associated muscles, generating a condition favorable for an alternative differentiation route towards dermal tissues.

The phenotype of the *Pax3Pr-Lbx1* transgenics indicates that *Lbx1* activity in the somites is context dependent. At limb and neck levels it is involved in the production of muscle progenitors ^29^; in trunk somites, however, it seems to interfere with myotomal-sclerotomal interactions leading to distal rib fusions. If the same is true in turtles, the latter activity could be involved in the formation of the peripheral element surrounding the carapace. Such *Lbx1* context-dependent activity would thus allow participation of the hypaxial somitic edge of trunk somites in carapace development without compromising limb muscle formation at forelimb and hindlimb levels.

It has been proposed that the protein produced from the *Myf5* splice variant lacking exon 2 could act as a dominant negative molecule ^20^. Our data, however, does not appear to support this interpretation, as heterozygous embryos for any of the *Myf5^turtle^* alleles are phenotypically normal in both skeletal and molecular patterns. Sequence analyses indicate that the frameshift resulting from skipping exon 2 still leaves a significant region of the transcriptional activating domain of the protein ^22^, and it is therefore possible that it keeps at least some of the normal activity of the molecule. Indeed, the phenotype of *Myf5^turtle^* embryos differs from that described for *Myf5* null mutants ^30^. Remarkably, the frameshift resulting from skipping exon 2 does not lead to an early stop codon but generates a 74 amino acid tail. This tail could influence Myf5 activity in different ways, from having no impact, to reducing its activity or even leading to the acquisition of novel functions. Running this tail through a variety of databases failed to identify similarities with known proteins or the presence of any known functional domain that could help to clarify a possible role for this part of the protein. The *Myf5* expression pattern in *Myf5^turtle^* embryos indicates that myotomal formation is largely conserved in these embryos. As myogenin has been reported to be under *Myf5* control ^31,32^, the reduced myogenin expression in *Myf5^turtle^* embryos might indicate decreased transcriptional activity of the Myf5^Δex2^ or Myf5^Δex2+Tr-ex3^ proteins. The *Myf6* expression pattern in *Myf5^turtle^* embryos requires a more complex explanation. The strongly reduced expression in the hypaxial myotome at E10.5 could also result from an eventual decrease in Myf5^Δex2^ transcriptional activity, were Myf5 required for *Myf6* activation. However, the premature *Myf6* activation in the hypaxial myotome of trunk somites requires either the loss of a Myf5-dependent repression of hypaxial *Myf6* expression or a gain of function activity in this embryonic region derived from Myf5^Δex2^ expression. Changes in the hypaxial dermomyotomal lip of *Myf5^turtle^* embryos are indeed observed in the form of premature de-epithelization. Interestingly, changes in the dermomyotome are not restricted to the hypaxial lip, as abnormal patterns were also observed in the anterior and posterior borders of this somitic compartment as illustrated by *Paraxis* expression. It will be interesting to examine whether the same is observed in turtle embryos.

The finding that the dermomyotome is affected in *Myf5^turtle^* embryos indicates the existence of a reverse flow of information from the myotome to the dermomyotome, which suggests the possibility that the extra 74 amino acid tail provides the Myf5^Δex2^ protein with an additional function promoting the production of signals affecting the overlying dermomyotome. Detailed analysis of the biochemical characteristics of Myf5^Δex2^ proteins will be required to clarify whether this is indeed the case.

## MATERIALS AND METHODS

### Mice and embryos

*Myf5^Tr-ex3^*, *Myf5^Δex2^* and *Myf5^Δex2+Tr-ex3^* mice were generated by CRISPR/Cas9 on the FVB/J background. In all cases the gRNAs were generated by hybridizing in vitro specific Alt-R® scRNAs with Alt-R® tracrRNA (both from IDT) and incubating them with Cas9 protein to generate an active ribonucleoprotein (RNP) complex. RNPs containing 0.61 μM of each hybridized gRNA and 50 ng/μl of Cas9, together with 30 ng/μl of a single-stranded (ss)DNA replacement oligonucleotide were microinjected into the pronucleus of fertilized oocytes and transferred into pseudopregnant females according to standard procedures ^33^. To produce both the *Myf5^Tr-ex3^* and the *Myf5^Δex2^* alleles we injected RNPs containing two gRNAs targeting the border of the sequence to be modified and one ssDNA oligo to promote the edition; the sequences targeted by the gRNAs and of the ssDNAs are shown in table 1. Genotyping of mice, fetuses and embryos carrying this allele was done by PCR using the oligonucleotides specified in table 1. Trans-heterozygous mice and embryos for the two *Myf5* alleles (*^Δex2/Tr-ex3^*) were generated by crossing homozygous mice for the *Myf5^Tr-ex3^* and the *Myf5^Δex2^* alleles. Finally, to generate the *Myf5^Δex2+Tr-ex3^* mice we used the strategy described above to remove the exon 2 but injecting oocytes recovered from homozygous *Myf5^Tr-ex3^* matings. In all cases the edited alleles were confirmed by direct sequencing of the target region.

**Table 1.**
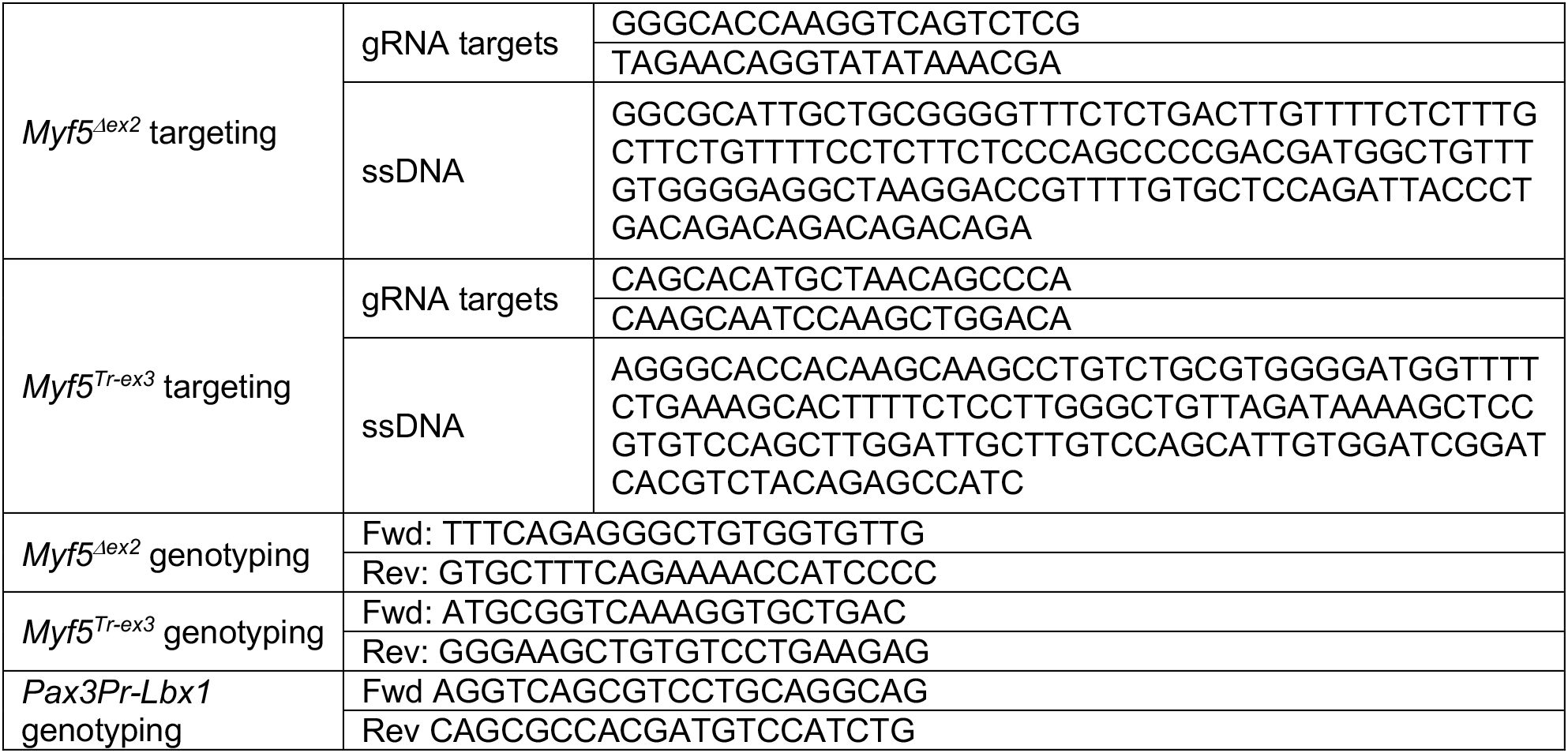

To generate *Pax3Pr-Lbx1* transgenic mice the mouse *Lbx1* cDNA (IMAGE clone 8733926) was cloned downstream of the hypaxial enhancer of the *Pax3* gene ^25^ and upstream of the SV40 polyadenylation signal. The construct was liberated from vector sequences, gel purified and microinjected into the pronucleus of fertilized FVB/J oocytes. Transgenic embryos (E10.5) and fetuses (E18.5) were identified by PCR using primers indicated in table 1.

All animal procedures were performed in accordance with Portuguese (Portaria 1005/92) and European (directive 2010/63/EU) legislations and guidance on animal use in bioscience research. The project was reviewed and approved by the Ethics Committee of “Instituto Gulbenkian de Ciência” and by the Portuguese National Entity “Direcção Geral de Alimentação Veterinária” (license reference: 014308).

### Whole mount *in situ* hybridization

Whole mount *in situ* hybridization was performed as described ^34^ using *in vitro* transcribed digoxigenin-labelled antisense RNA probes. Briefly, embryos were dissected out in Dulbecco’s Phosphate Buffered Saline (PBS) (1.8 mM KH2PO4, 2.7 mM KCl, 10 mM Na_2_HPO_4_, 137 mM NaCl) and fixed with 4% paraformaldehyde (PFA) made in PBS at 4°C overnight. Embryos were washed in PBT (PBS containing 0.1% Tween-20), dehydrated with methanol and rehydrated with PBT. They were then bleached in 6% hydrogen peroxide, treated with proteinase K (10 μg/ml in PBT), the reaction was stopped with glycine (2 mg/ml in PBT) and embryos were postfixed with 4% PFA, 0.2% glutaraldehyde. Hybridization was performed at 65°C overnight in hybridization solution [50% formamide, 1.3 x SSC (3M NaCl, 300 mM sodium citrate, pH 5.5), 5 mM EDTA, 0.2% Tween 20, 50 μg/ml yeast tRNA, 100 μg/ml heparin] containing the RNA probe, followed by three washes at 65°C in hybridization solution without the RNA probe, tRNA and heparin. Embryos were then washed in TBST (25 mM Tris.HCl, pH 8.0, 140 mM NaCl, 2.7 mM KCl, 0.1% Tween 20), equilibrated with MABT (100 mM Maleic acid, 150 mM NaCl, 0.1% Tween-20, pH 7.5), blocked with MABT/Block [MABT containing 1% blocking reagent (Roche #11096176001) and 10% sheep serum] and incubated with a 1:2000 dilution of alkaline phosphatase-conjugated anti-digoxigenin antibody (Roche Cat# 11093274910) in MABT/Block at 4°C overnight. Embryos were washed extensively with MABT at room temperature, equilibrated in NTMT (100 mM Tris HCl, pH 9.5, 50 mM MgCl_2_, 100 mM NaCl, 0.1% Tween-20) and developed at room temperature with NBT/BCIP (Roche #11681451001) diluted in NTMT. Reactions were stopped with PBT, the embryos were fixed with 4% PFA and stored in PBT. At least 3 embryos were stained per probe and genotype, showing highly reproducible patterns.

To section the stained embryos, they were embedded in gelatin/albumin [0.45 % gelatin, 270 g/l bovine serum albumin, 180 g/l sucrose in PBS, jellified with 1.75 % glutaraldehyde], sectioned with a vibratome at 35 μm and mounted with an aqueous mounting solution (Aquatex, Merck).

### Skeletal staining

E18.5 fetuses were stained as previously described ^35^. Briefly, fetuses were skinned, eviscerated, and fixed in 100% ethanol for 24 hours. They were then incubated at room temperature for 12 hours in 150 mg/l of alcian blue 8 GX made in 20% acetic acid 80% ethanol and post fixed in 100% ethanol for 24 hours. They were then cleared in 2% KOH for 8 hours, stained in 50 mg/l alizarin red S in 2% KOH for 3 hours, and further cleared overnight in 2% KOH. The process was finished by replacing the KOH with 25% glycerol in H2O and stored in the same solution.

### Histological analysis

E18.5 fetuses were fixed in Bouin’s fixative for 2 days at room temperature, dehydrated extensively with repeated changes of 100% ethanol for 5 days, embedded in paraffin, sectioned at 10 μm and stained with Masson’s trichrome according to standard methods.

## ACKNOWLEDGEMENTS

We would like to thank the members of the Mallo lab, especially Ana Casaca and Anastasiia Lozovska for practical guidance during several steps of this project, the IGC mouse facility for their help with animal housing and histopathology facility for helping with histological procedures. This project was funded by grant PTDC/BIA-BID/30254/2017 (from FCT, Portugal) to MM and the research infrastructure Congento, project LISBOA-01-0145-FEDER-022170.

